# DNAharvester: A Nextflow Pipeline for Analysing Highly Degraded DNA from Ancient and Historical Specimens

**DOI:** 10.64898/2026.04.20.719564

**Authors:** Bilal Sharif, Verena E. Kutschera, Nikolay Oskolkov, Benjamin Guinet, Edana Lord, J. Camilo Chacón-Duque, Jonas Oppenheimer, Tom van der Valk, David Díez-del-Molino, Peter D. Heintzman, Love Dalén

## Abstract

Ancient DNA (aDNA) research has advanced rapidly with the development of high-throughput sequencing, enabling genome-wide analyses of large collections of prehistoric specimens. However, analysing palaeontological and archaeological material with highly degraded DNA constitutes a major bioinformatic challenge. DNA from such samples is characterised by short fragment lengths, low endogenous content, post-mortem damage, and cross-species contamination, which can increase spurious mapping and reference bias, affecting downstream population genetic inferences. We present DNAharvester, a modular and reproducible pipeline designed specifically for processing highly degraded DNA from ancient and historical specimens. DNAharvester integrates metagenomic filtering, competitive mapping, adaptive aligner selection (incorporating *BWA-aln, BWA-mem*, and *Bowtie2*), and systematic evaluation of reference bias and spurious mapping. By incorporating flexible mapping and filtering strategies, the pipeline can be adapted to varying sample preservation, focusing on maximising authentic data recovery. DNAharvester features subworkflows for iterative assembly of mitogenomes, identification of genomic repeats and CpG sites, taxonomic classification, microbial/pathogen screening, genetic sex determination, and variant calling. To accommodate varying sequencing depths, the pipeline supports diploid variant calling, genotype likelihood estimation, and pseudo-haploid random allele calling. Implemented in Nextflow, DNAharvester provides a highly scalable, containerised framework that enhances reproducibility, portability, and robustness in aDNA analyses. We validated the pipeline using simulated and empirical datasets, demonstrating its ability to systematically mitigate complex background contamination while preserving authentic genomic signals. By streamlining complex bioinformatic tasks through simple configuration files, DNAharvester establishes a standardised approach for analysing aDNA datasets and makes genomic analyses of ancient remains accessible to the broader research community.

## 1. Background

The rapid development and decreasing cost of high-throughput sequencing has fundamentally transformed our ability to reconstruct the evolutionary history of species. As sequencing large quantities of DNA molecules has become increasingly feasible, ancient DNA (aDNA) research has shifted from analysing short fragments of organellar DNA to generating large genome-wide datasets comprising hundreds of samples from past populations [1]. These datasets allow researchers to investigate evolutionary events of populations, such as migration, admixture, expansion and decline, and extinction, that are often beyond the realm of morphological analyses [2,3]. To date, most aDNA studies have focused on material from the Late Pleistocene and Holocene, typically within the last ∼50,000 years [4]. This temporal skew largely reflects the limit of radiocarbon dating, the relative abundance of samples, and DNA preservation constraints, whereby cold environments and relatively recent samples are more likely to retain usable DNA [1,5]. However, recent methodological advances are extending the temporal and preservational limits of genome recovery. With the advent of deep-time palaeogenomics, genomic data have been retrieved from materials more than a million years old [6,7].

However, pushing these boundaries presents substantial biological and computational challenges. From the moment of death, through to deposition, field sampling, and subsequent laboratory processing, the sample continuously accumulates exogenous DNA from microorganisms and other sources, such as human handling. The small fraction of surviving original DNA, the endogenous fraction, is often vastly outnumbered by a complex exogenous mixture of modern and ancient contaminants. Furthermore, these endogenous DNA molecules are highly fragmented and carry post-mortem chemical damage, particularly cytosine deamination [8]. Moreover, the exogenous DNA present in the sample can be as ancient as the host organism itself, meaning these contaminants have been subjected to the same fragmentation and chemical damage as endogenous molecules. This creates a severe bioinformatic challenge, as ancient endogenous and exogenous DNA molecules cannot be distinguished based on standard authentication criteria.

These challenges are compounded, especially in the case of extinct species and/or individuals from genomic deep-time (>126,000 years old), where researchers must map their aDNA sequences to divergent reference genomes. Standard bioinformatic tools developed for high-quality modern genomes do not perform optimally under these conditions. DNA alignment performance varies strongly with read length, sample-to-reference sequence divergence, and post-mortem damage level [9], and no single mapping strategy performs equally well across all preservation states. For example, the most widely used algorithm in aDNA studies, *bwa-backtrack*, performs better on ultrashort reads (∼30–70 bp), whereas *bwa-mem* is optimised for longer fragments (>70 bp) [9]. Using a single, fixed mapping strategy can therefore lead to data loss, spurious alignments, and/or reference bias, which each occur to different extents among samples. Such biases can affect downstream population genetic inferences.

To mitigate these issues, several complementary tools and strategies have been proposed. One approach is competitive mapping, in which sequencing reads are aligned against a concatenated reference genome comprising the target organism and a decoy genome (e.g., human when targeting a faunal taxon) [10]. Contaminant reads are at least equally likely to map to the decoy, thereby reducing their mapping quality without losing an appreciable proportion of endogenous DNA [10]. However, this approach becomes computationally demanding when contamination is taxonomically complex. Another strategy involves filtering data through metagenomic classification tools such as Kraken 2 prior to genome mapping [11], which can efficiently remove microbial contaminants before they reach the mapping stage. In addition, the recently developed AMBER tool [9] enables systematic evaluation of mapping quality by assessing mismatch rates across read length bins, allowing the detection of reference bias and spurious alignments.

Despite the efficacy of these individual tools and strategies, integrating them into a coherent and reproducible workflow has not yet been done. Because palaeogenomic data processing is highly complex and study-specific, researchers frequently rely on ad hoc scripts to chain disparate software together. This methodological fragmentation not only makes data processing cumbersome but also severely hinders reproducibility, making it difficult to independently replicate findings or deploy analyses across different computing environments.

To fill this gap, we introduce DNAharvester, a Nextflow pipeline that provides a robust, containerised solution specifically designed for highly degraded ancient and historical DNA datasets. DNAharvester compiles state-of-the-art software and techniques for processing such samples, including metagenomic filtering prior to mapping, competitive mapping, and AMBER-based mapping evaluation, into a single, reproducible pipeline. We detail the pipeline’s modular architecture and benchmark its performance across a gradient of simulated scenarios and empirical datasets, demonstrating that authentic endogenous DNA can be reliably and consistently recovered from the most challenging material.

## 2. Pipeline Workflows and Data Processing

The pipeline is designed with a modular architecture comprising different subworkflows that can be toggled on or off. By default, DNAharvester runs a set of subworkflows (Fig. 1) that checks the validity of input data/parameters (input_check); performs quality control of raw paired-/single-end sequencing reads (raw_fastq_qc); removes adapter sequences and merges paired-end reads (fastq_processing); aligns the processed reads to the reference genome (mapping) or a competitive reference (competitive-mapping); performs quality control of raw alignments (raw_bam_qc); filters the alignments for quality and duplicates (bam_processing); performs quality control of the final alignments (processed_bam_qc); and compiles statistics of read processing and alignment at different steps of the processes (mapping metrics). Additionally, DNAharvester includes specialised subworkflows for the identification of repetitive elements and CpG sites in the reference genome (repeat_cpg_identification); variant calling using hard-called genotypes, genotype likelihoods, and pseudo-haploid calls by random allele sampling (variant_calling); iterative assembly of mitogenomes (iterative_assembly); taxonomic classification of the samples (taxonomic_classification); microbial/pathogen screening (microbial_screening); and genetic sex determination (sexing) (Fig. 1). These subworkflows are described in the sections below.

**Fig. 1.**
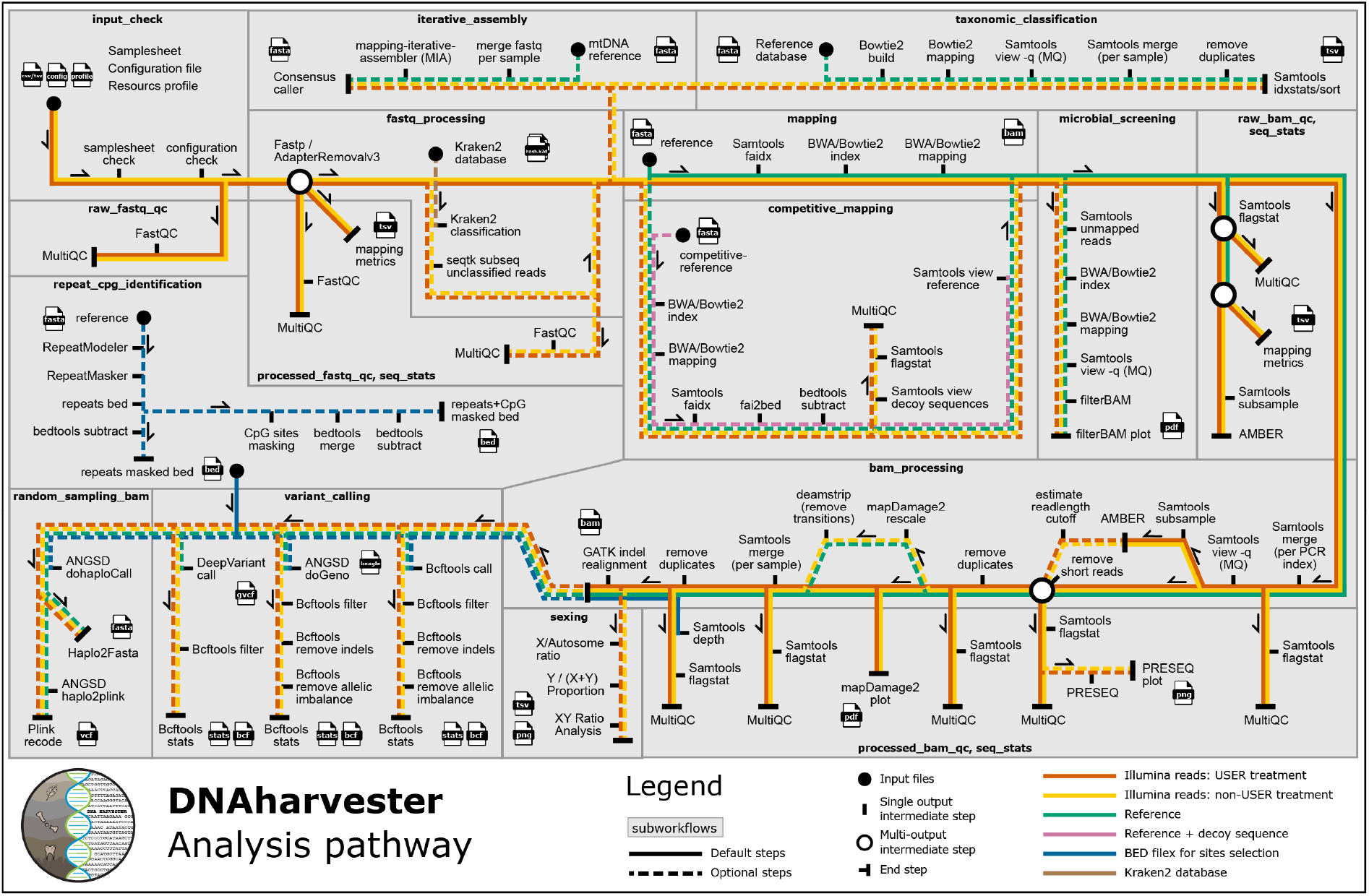
Schematic overview of the DNAharvester analysis workflow. The flowchart shows the different subworkflows (separated by grey outlines) and the direction of data flow among them. Line colours indicate different data types.

### 2.1. Input check

The pipeline is configured using a sample sheet and a configuration file. The sample sheet links sample IDs with metadata, including the sample type, library type, sequencing library ID, sequencing platform, flow cell ID, sequencing lane number, and paths to the input raw data in fastq format. The pipeline accommodates diverse experimental setups, supporting multiple libraries per sample and multiple lanes/fastq files per library. It is compatible with both single- and double-stranded DNA library datasets. Before starting a run, the pipeline performs a validation check of the sample sheet and configuration settings. This ensures that all necessary files are present and that no conflicting parameters have been set in the configuration file.

### 2.2. Fastq processing

In the fastq processing subworkflows, sequencing adapters are removed and paired-end reads can optionally be merged using Fastp [12] by default, or alternatively using AdapterRemoval3 [13]. If enabled, remaining unmerged reads can also be retained. When Fastp is used, a novel custom tool named “AdaptClean” [14] can be enabled for single-end and unmerged paired-end reads to remove residual adapter sequences shorter than 5 bp. This is important because Fastp requires a minimum overlap of 5 bp to detect adapter sequences and so will not trim adapter sequences from library inserts that approach the read length (see Supplementary Text heading S1). Additionally, the shortest reads can be filtered by setting a minimum read length threshold in the configuration file. Alternatively, this parameter can be set to ‘auto’; in this mode, reads as short as 20 bp are initially retained, and a library-specific threshold is estimated after mapping using AMBER [9] (see BAM Processing). As an optional pre-mapping filtering step, processed reads can be classified using Kraken 2’s *k*-mer classification [15] against a user-provided database (e.g., the Genome Taxonomy Database of bacteria and archaea (GTDB) [16]), whereby only unclassified reads are retained. This step reduces the spurious mapping of exogenous contaminants, thereby improving overall mapping efficiency [11]. Quality control of both the raw and processed fastq reads is performed using FastQC [17] and MultiQC [18].

### 2.3. Mapping

DNAharvester supports the alignment of processed reads to a reference genome using one of four mapping strategies: *bwa-aln, bwa-mem*, a hybrid *bwa-aln-mem* approach, or *bowtie2* [19–21]. *Bwa-aln* is recommended for short ancient DNA reads (< ∼70 bp), while *bwa-mem* works best for relatively longer reads (> ∼70 bp) from both ancient/historical and modern samples (see Supplementary Text heading S2; [9,19]). *Bowtie2* is recommended when mapping against a large database due to its lower memory requirements and faster processing time, but is more prone to short read reference bias than *bwa-aln* [9]. For samples containing both short and longer reads, we include a novel hybrid approach, *bwa-aln-mem*, in which short reads are aligned using *bwa-aln* and reads longer than a user-specified threshold (default: 70 bp, configurable) are aligned with *bwa-mem* (Supplementary Text heading S2).

Mapping parameters are automatically adjusted based on sample type. By default, for ancient samples, *bwa-aln* uses “-l 1024 -n 0.01 -o 2”, and *bwa-mem* uses “-k 19 -r 2.5 -L 15” [22,23]. For modern samples, default parameters are used for both *bwa-aln* and *bwa-mem*. For both ancient and modern samples, *bwa-mem*’s soft-clipping behaviour, where the aligner trims the ends of reads rather than forcing a full alignment, is disabled by increasing the clipping penalty (-L 15). This ensures that the full read length is considered during alignment, which helps prevent short, spurious partial matches. However, soft-clipping can be useful when mapping longer reads to a divergent reference genome, and can be enabled in the configuration file. *Bowtie2* defaults to “--sensitive” for both ancient and modern samples [24]. All mapping parameters can be customised in the configuration file.

To further reduce spurious mapping of non-endogenous reads in ancient samples, the pipeline supports competitive mapping. In this mode, reads are aligned against a concatenated reference composed of the target species genome and decoy sequences (e.g., human and microbial genomes). The resulting BAM files are then split, retaining only reads that map uniquely to the target genome. This approach has been shown to significantly reduce spurious mapping, especially in poorly preserved ancient and historical samples with a high contamination load [10].

### 2.4. BAM processing

In the BAM processing subworkflows, BAM files are first merged per library ID, and reads with low mapping quality (default: MQ<25) are filtered out. The merged BAM files are then analysed with AMBER [9] to evaluate potential mapping issues, including reference bias, spurious alignments, and damage patterns. Here, if the read length parameter was set to ‘auto’ during fastq processing, the pipeline estimates an optimal minimum read-length cutoff based on the mismatch rate for each read length (see Supplementary Text heading S3), and reads shorter than the cutoff are filtered out. PCR duplicates are then removed using a custom Python script [25] that considers duplicates as having identical mapping coordinates. The resulting BAM files are then assessed for post-mortem damage profiles using MapDamage2 [26]. To address aDNA damage in non-UDG-treated libraries (where uracil-DNA glycosylase (UDG) [27] has not been applied during library preparation to remove uracils resulting from deaminated cytosines), we implemented two options. The first is rescaling the BAM files using MapDamage2’s “--rescale” parameter [26], which models cytosine deamination damage using a Bayesian framework and reduces base quality scores to 0 for likely damaged bases. Alternatively, users can apply deamstrip, a novel custom tool that mitigates deamination in a strand-specific manner. For single-stranded libraries, it removes C-to-T substitutions on the forward strand and G-to-A substitutions on the reverse strand. For double-stranded libraries, it removes both substitution types. This is useful when a limited number of mapped reads prevents MapDamage2 from building the Bayesian model for damage estimation. BAM files are then merged per sample ID, and a second round of deduplication is performed to remove any remaining duplicate artifacts. Finally, an optional indel realignment step can be performed using GATK3 [28]. While this step can be disabled in the configuration file to reduce computational time, it is highly recommended when mapping to divergent reference genomes, as it effectively resolves misalignment artifacts, such as false SNPs near the ends of reads (see Supplementary Text heading S4).

Quality control of the raw, intermediate, and fully processed BAM files is performed using SAMtools flagstat [29] and MultiQC [18] after each major step, including MQ filtering, short read removal, BAM file merging, and deduplication. Additionally, library complexity can be estimated using Preseq’s lc_extrap [30] to predict the number of uniquely mapped reads expected with further sequencing for each library. A plot visualising the expected number of uniquely mapped reads versus total mapped reads is generated using a novel custom Python script. The mean depth of coverage is also calculated for each sample using the fully processed, deduplicated BAM files. By default, this calculation considers all positions across the reference genome. However, a BED file can be provided to restrict the calculation to specific regions of interest, such as capture targets or non-repetitive sites generated by the genomic repeats and CpG sites identification subworkflow (described below).

### 2.5. Identification of genomic repeats and CpG sites

This subworkflow processes the target reference genome to identify repetitive elements and CG dinucleotides (CpG sites). First, repetitive and low complexity sequences are identified using RepeatModeler [31] and RepeatMasker [32]. These regions are prone to spurious alignments, and masking them from the reference genome greatly reduces false positives during the variant calling step. The subworkflow creates BED files containing repeat regions and their respective masked counterparts. Second, a custom Python script [33] locates all CpG sites across the genome. CpG sites require special attention due to the nature of cytosine deamination damage [27]. Unlike cytosines at non-CpG sites, cytosines at CpG sites are often methylated by epigenetic modification. Whereas cytosine deaminates to uracil, a methylated cytosine deaminates to thymine, which will not be removed by UDG treatment. Consequently, even in UDG-treated libraries, CpG sites retain deamination damage that can bias the downstream variant calling step. However, the surviving damage pattern at CpG sites can be leveraged to authenticate the UDG-treated ancient DNA [34], also implemented in AMBER [9]. We note that this does not seem to apply to animal mitochondrial genomes. The subworkflow generates BED files for CpG sites and their respective masked counterparts, as well as a combined file with both repeats and CpG sites masked. These BED files allow users to restrict downstream analyses, such as depth of coverage calculations and variant calling, to non-repetitive and non-CpG regions.

### 2.6. Variant calling

DNAharvester includes three distinct variant calling options, tailored to accommodate datasets ranging from low to high sequencing depth. For high-coverage samples (typically >7×), the user can select BCFtools (mpileup/call) [29] or DeepVariant [35] for variant hard-calls. The resulting variants are then filtered based on minimum and maximum depth and quality, with additional filters for removing indels and allelic imbalance for BCFtools. For medium-coverage samples (approx. 3–7×), the user can select ANGSD to call variants in a genotype likelihood framework [36]. For low-coverage samples (<3×), ANGSD can be selected to generate pseudo-haploid calls by randomly sampling a single allele per site (-doHaploCall 1) [36]. All of the above variant calling options can be restricted to user-defined target regions or specifically configured to exclude the repetitive regions and CpG sites identified in the previous subworkflow.

### 2.7. Iterative assembly of mitogenomes and small nuclear regions

The iterative assembly subworkflow incorporates the mapping-iterative-assembler (MIA) [37] to reconstruct mitogenomes and small nuclear regions. First, all fastq files from the fastq processing step above are merged per sample ID. MIA then aligns these reads to a user-provided seed reference genome, calls a consensus sequence, and utilises this newly generated consensus as the reference for subsequent alignment rounds. This iterative process continues until sequence convergence is achieved. The final output from MIA is then processed using a custom Python script [38] to generate the consensus sequence in fasta format, applying minimum thresholds for read depth (default: 3x), base quality (default: 0), and allele consensus agreement (default: 66%).

### 2.8. Taxonomic classification

The taxonomic classification subworkflow is designed to identify samples of unknown, dubious, or mixed identity. First, the processed fastq files are aligned against a user-provided reference database (e.g., the RefSeq mitochondrial database) using *bowtie2*, with up to 10 multi-mapping reads retained by default. Alignments with low mapping quality (default: MQ<25) are removed. Subsequently, the filtered BAM files are merged per sample ID, and PCR duplicates are removed. Finally, SAMtools idxstats is run on the processed BAM files to tabulate mapped reads and identify the reference sequence with the highest read count, thereby inferring the most likely taxonomic assignment(s).

### 2.9. Microbial screening

The microbial screening subworkflow processes unmapped reads from the primary mapping step to identify potential microbes or pathogens present in the samples. The unmapped reads are aligned against a user-provided microbial reference database, and alignments with low mapping quality (default: MQ<1) are filtered out. Subsequently, the resulting BAM files are merged per sample ID, and PCR duplicates are removed. FilterBAM [39] is then used to collect mapping metrics. Finally, a custom Python script generates diagnostic plots for candidate microbes, visualising key authentication and abundance metrics such as the number of mapped reads, average mapping quality, edit distance, average nucleotide identity, coverage evenness, sequence identity percentage, read length distribution, and aDNA damage patterns. It is important to note that this subworkflow is designed to provide a rapid initial screening prediction based on direct alignment. For a comprehensive metagenomic analysis, samples should be subsequently processed using dedicated metagenomic pipelines (e.g., aMeta [40] or FALCON2 [41]).

### 2.10. Genetic sex determination

The genetic sex determination subworkflow infers the biological sex of samples with XY chromosomal systems using read count ratios extracted from processed BAM files, and can easily be used for other systems, such as ZW. Depending on which target chromosome names (X, Y, and/or autosomes) are specified in the configuration file, the pipeline dynamically adapts its approach. If only the names of the X chromosome and autosomes are provided, genetic sex is determined by comparing the length-normalised read depth of the X chromosome to the overall autosomal depth. By default, a ratio above 0.8 is considered female (XX), and a ratio below 0.6 is considered male (XY). This method also generates a visualisation of relative chromosome ploidy to highlight the expected diploid or haploid coverage of the X chromosome. If both the X and Y chromosome names are specified, the pipeline calculates the fraction of sequences aligned to the Y chromosome relative to the total number of sequences aligned to either sex chromosome (Y / (X+Y)) [42]. Due to potential variations in Y chromosome assembly quality across different reference genomes, the pipeline provides this proportion for user interpretation rather than providing an automated call. Finally, if all three parameters are provided, the pipeline generates a scatter plot comparing the Rx (X chromosome read count relative to total autosomal read count) and Ry (Y chromosome read count relative to total autosomal read count) across samples [43].

### 2.11. Output

The pipeline outputs all generated files to a user-defined results directory, structured according to the executed subworkflows. This includes processed fastq files, raw and processed BAM files, and comprehensive mapping metrics aggregated into tsv files. These metrics encompass counts for raw reads, adapter-filtered reads, total mapped reads, MQ-filtered reads, endogenous DNA percentage (defined as MQ-filtered reads / raw reads), uniquely mapped reads, and library complexity (defined as uniquely mapped reads / MQ-filtered reads), along with read length statistics (minimum, maximum, mean, and median). To facilitate granular quality control, mapping metrics are generated at two distinct stages: first, after merging BAM files per sequencing library, and second, after merging all libraries per sample.

## 3. Benchmarking and Discussion

### 3.1. Simulated scenario datasets

To evaluate the performance of the fastq processing, mapping, and BAM processing subworkflows, we simulated five samples (three libraries per sample) with varying levels of endogenous content and contamination, as well as distinct read length distributions (Table 1, Fig. 2). Because the true origin of each read is known, these simulated datasets allow for a direct assessment of spurious alignments, contaminant removal, and endogenous DNA recovery [9]. Raw reads were simulated using Gargammel [44], with (1) chromosome 1 of the African elephant (*Loxodonta africana*) reference genome (GenBank: NC_087342.1) as the endogenous target, (2) human (*Homo sapiens*) chromosome 1 (GenBank: NC_000001.11) as a modern contaminant, (3) the bubonic plague bacterium (*Yersinia pestis*) genome (RefSeq: GCF_900460465.1) for pathogen detection, and (4) microbial genomes from the GTDB (release 226) [16] as environmental noise. To test the iterative assembly of the mitogenome and taxonomic classification subworkflows, we also included a varying number of simulated woolly mammoth (*Mammuthus primigenius*) mitogenome reads (derived from GenBank accession NC_007596.2) (Tables 2-3). These simulated samples represent a degradation gradient ranging from modern, high-quality DNA to highly degraded aDNA samples. This diverse dataset enabled robust testing of the pipeline’s ability to successfully merge multiple libraries, mitigate spurious alignments, and accurately assign reads to their true genomic origin. All scripts used to generate this simulated dataset and the subsequent DNAharvester configuration files are provided on GitHub.

**Table 1:**
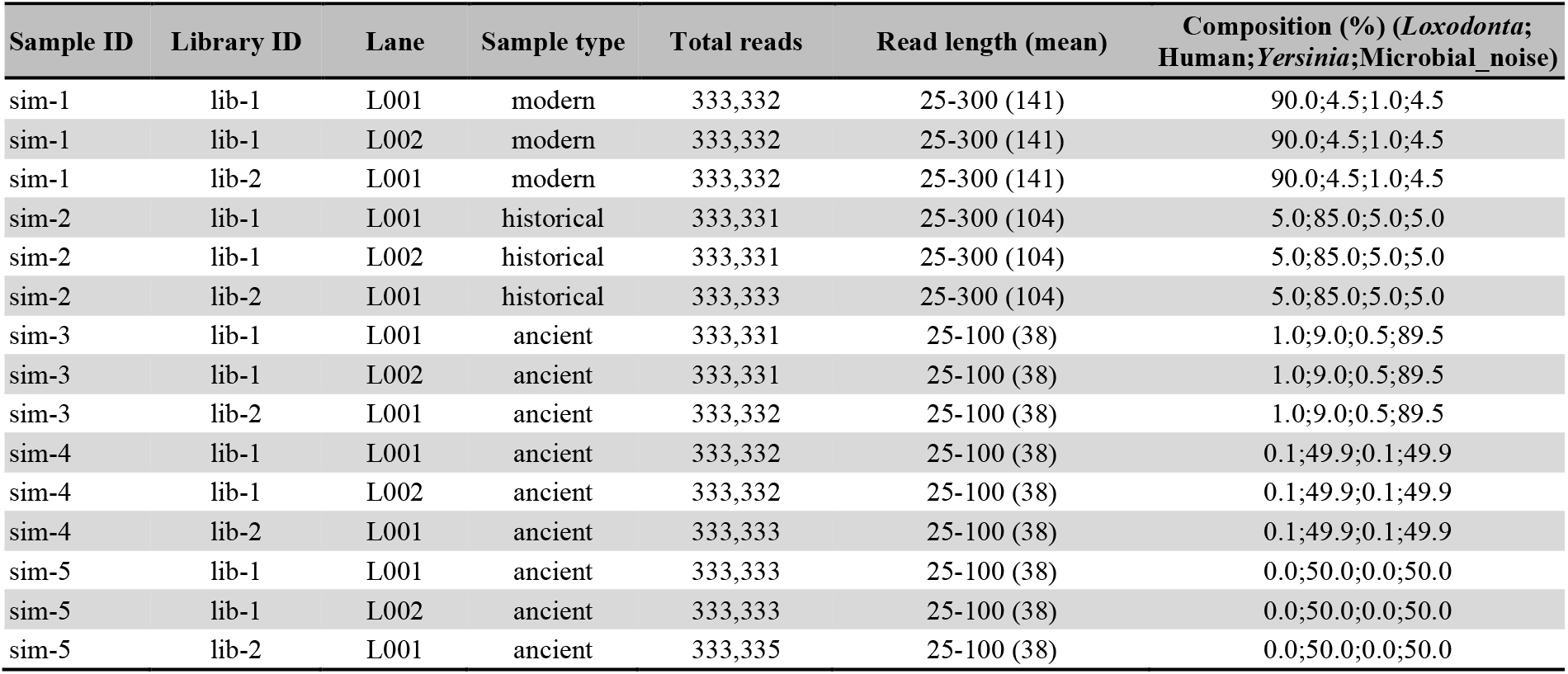
Simulated sample dataset specifications. Summary of the five simulated sample profiles (sim-1–sim-5) comprising varying proportions of endogenous (*Loxodonta africana*), contaminant (*Homo sapiens*), pathogen (*Yersinia pestis*), and microbial noise (GTDB)

| Sample ID | Library ID | Lane | Sample type | Total reads | Read length (mean) | Composition (%) (*Loxodonta*;<br>Human;*Yersinia*;Microbial_noise) |
| --- | --- | --- | --- | --- | --- | --- |
| sim-1 | lib-1 | L001 | modern | 333,332 | 25-300 (141) | 90.0;4.5;1.0;4.5 |
| sim-1 | lib-1 | L002 | modern | 333,332 | 25-300 (141) | 90.0;4.5;1.0;4.5 |
| sim-1 | lib-2 | L001 | modern | 333,332 | 25-300 (141) | 90.0;4.5;1.0;4.5 |
| sim-2 | lib-1 | L001 | historical | 333,331 | 25-300 (104) | 5.0;85.0;5.0;5.0 |
| sim-2 | lib-1 | L002 | historical | 333,331 | 25-300 (104) | 5.0;85.0;5.0;5.0 |
| sim-2 | lib-2 | L001 | historical | 333,333 | 25-300 (104) | 5.0;85.0;5.0;5.0 |
| sim-3 | lib-1 | L001 | ancient | 333,331 | 25-100 (38) | 1.0;9.0;0.5;89.5 |
| sim-3 | lib-1 | L002 | ancient | 333,331 | 25-100 (38) | 1.0;9.0;0.5;89.5 |
| sim-3 | lib-2 | L001 | ancient | 333,332 | 25-100 (38) | 1.0;9.0;0.5;89.5 |
| sim-4 | lib-1 | L001 | ancient | 333,332 | 25-100 (38) | 0.1;49.9;0.1;49.9 |
| sim-4 | lib-1 | L002 | ancient | 333,332 | 25-100 (38) | 0.1;49.9;0.1;49.9 |
| sim-4 | lib-2 | L001 | ancient | 333,333 | 25-100 (38) | 0.1;49.9;0.1;49.9 |
| sim-5 | lib-1 | L001 | ancient | 333,333 | 25-100 (38) | 0.0;50.0;0.0;50.0 |
| sim-5 | lib-1 | L002 | ancient | 333,333 | 25-100 (38) | 0.0;50.0;0.0;50.0 |
| sim-5 | lib-2 | L001 | ancient | 333,335 | 25-100 (38) | 0.0;50.0;0.0;50.0 |

**Table 2.** Summary of iterative assembly results. Accuracy and completeness of the reconstructed woolly mammoth (*Mammuthus primigenius*) mitogenomes, generated using DNAharvester’s iterative assembly subworkflow. Varying amounts of simulated woolly mammoth mitogenome reads were spiked into five simulated datasets (sim-1 to sim-5). The Asian elephant (*Elephas maximus*) mitogenome (GenBank: CM123153.1) was used as the seed reference.

| Sample ID | Reads added per sample | Reference length (bp) | Assembled (bp) | Completeness (%) | Errors (SNPs/Indels) | Accuracy (%) |
| --- | --- | --- | --- | --- | --- | --- |
| sim-1 | 300 | 16,770 | 1760 | 10.49 | 35 | 98.01 |
| sim-2 | 3,000 | 16,770 | 7275 | 43.38 | 42 | 99.42 |
| sim-3 | 6,000 | 16,770 | 16017 | 95.51 | 3 | 99.98 |
| sim-4 | 15,000 | 16,770 | 16070 | 95.83 | 0 | 100.00 |
| sim-5 | 30,000 | 16,770 | 16120 | 96.12 | 2 | 99.99 |

**Fig. 2.**
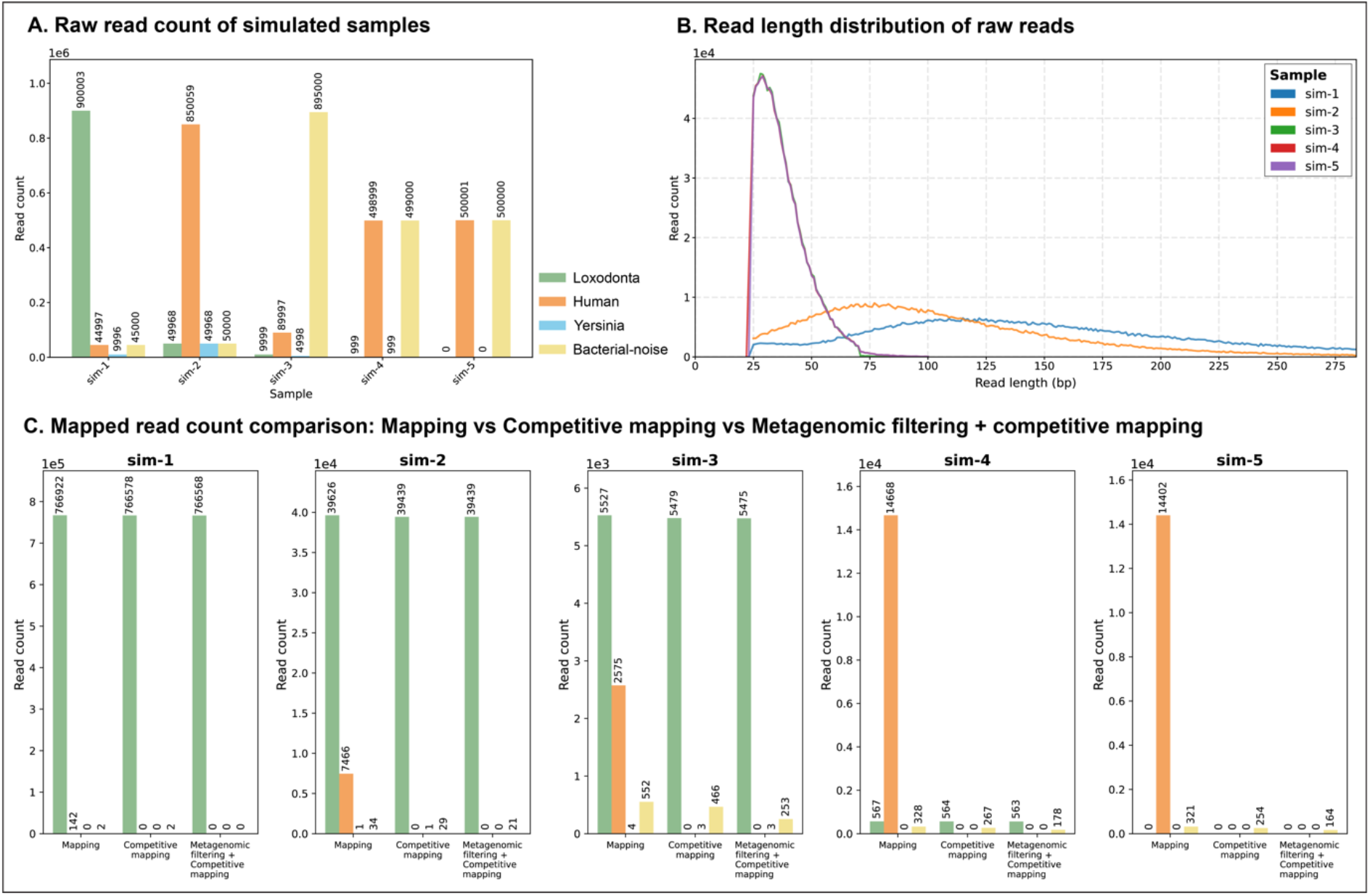
Characteristics and mapping evaluation of the simulated sample datasets. (A) Proportional composition of the raw simulated reads across the five samples (sim-1 to sim-5) (B) Read length distributions for the simulated datasets (C) Comparison of mapped read compositions resulting from three distinct processing strategies: standard mapping, competitive-mapping, and Kraken 2 filtering combined with competitive mapping.

### 3.2. Standard mapping, competitive mapping, and metagenomic filtering

DNAharvester was initially evaluated on the simulated sample datasets using standard default mapping parameters (*bwa-aln*, read merging enabled, minimum read length: 30 bp, mapping quality: 25). For the standard mapping baseline, reads were aligned solely to the Asian elephant (*Elephas maximus*) reference genome (GCF_024166365.1). To evaluate the competitive-mapping subworkflow, reads were aligned against a concatenated reference containing both the target *Elephas maximus* genome and the decoy *Homo sapiens* genome (GCF_000001405.40). For the metagenomic filtering step prior to mapping, the Kraken 2 database was built with default parameters except for the k-mer size set to 30 bp, and classified reads were filtered with a confidence threshold of 0.05.

Standard mapping performed well on the simulated modern sample (sim-1), recovering 88.25% of true *Loxodonta* reads (mentioned hereafter as recall rate), and 99.98% of mapped reads were true *Loxodonta* reads (hereafter referred to as precision) (Fig. 3). However, mapping precision steeply declined as read lengths decreased and contamination levels increased. In the simulated highly contaminated sample (sim-2), spurious alignments from human contaminant reads reduced the precision to 84.08%. Activating the competitive-mapping subworkflow effectively resolved the issue of human contamination (precision: 99.92%), while keeping the recall rate similar. In the case of a sample with both high human and microbial contamination, the combination of metagenomic filtering and competitive mapping performed best, increasing precision without substantially reducing the recall rate (Fig. 3). For example, in sim-3, precision increased from 92.11% (competitive-mapping) to 95.53% (metagenomic filtering plus competitive-mapping), while the recall rate remained almost the same. This precision improvement was even more pronounced in the highly degraded sim-4 sample, where precision increased from 67.87% to 75.98%, while recall experienced only a minor drop (from 70.85% to 70.73%). In sim-5, which contained no true endogenous DNA (serving as a negative control), the combined approach of competitive mapping and metagenomic filtering yielded the lowest number of spuriously mapped reads, thereby minimising false positives.

**Fig. 3.**
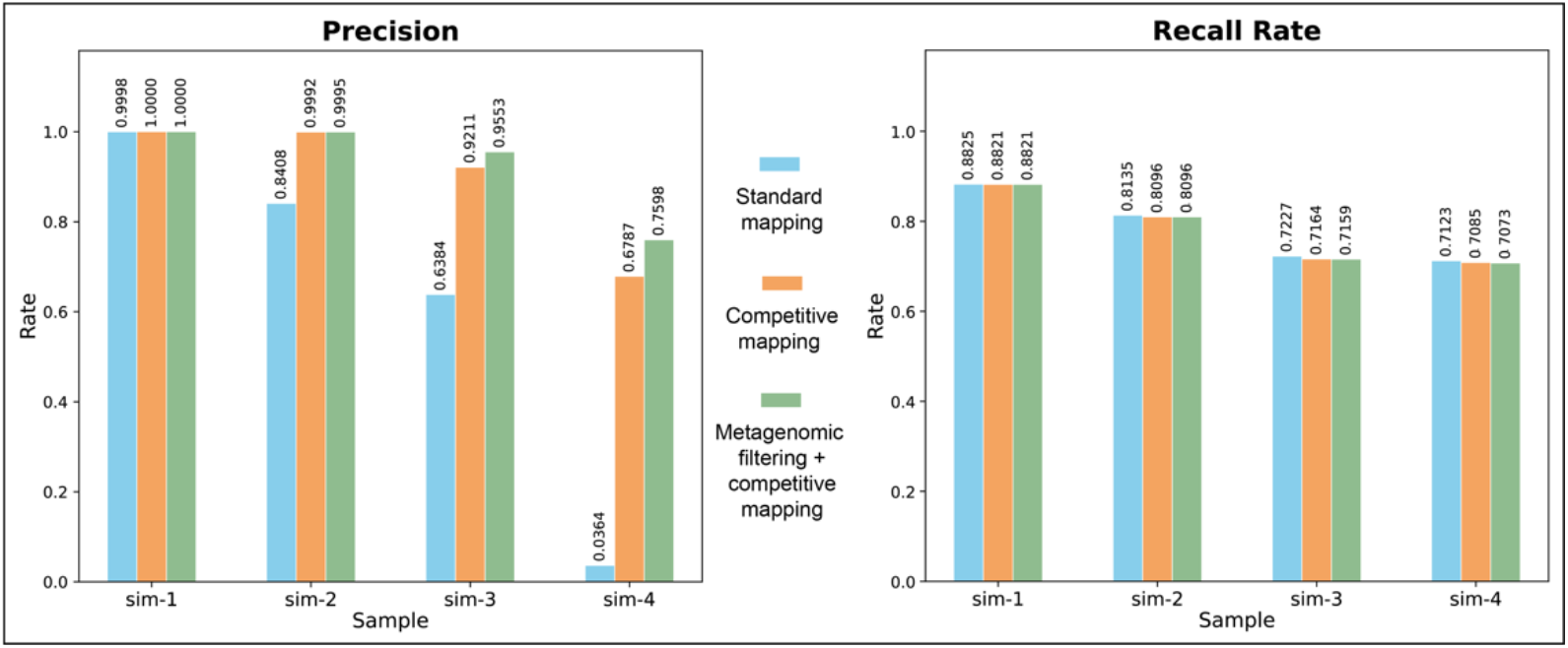
Precision and recall rate of different mapping strategies available in DNAharvester. (Left) Precision across the simulated datasets, defined as the fraction of all reads mapped to the target genome that are authentic *Loxodonta* reads (True Positives / [True Positives + False Positives]). (Right) Recall across the simulated datasets, defined as the fraction of all authentic *Loxodonta* reads present in the sample that the pipeline successfully recovered (True Positives / [True Positives + False Negatives]).

Together, these results demonstrate that standard mapping alone markedly underperforms at low-endogenous DNA conditions, where spurious alignments can easily dominate the dataset and artificially inflate coverage estimates. Competitive mapping provides a robust solution for substantially reducing host- and closely related-contamination, while metagenomic filtering offers the most efficient mechanism for removing distant microbial noise. The integration of these approaches within DNAharvester enables highly specific discrimination between authentic endogenous reads and complex contaminant sequences, which is strictly necessary in scenarios characterised by extremely low endogenous content.

### 3.3. Comparison of mapping tools

We further evaluated the performance of four mapping tools supported by the pipeline (*bwa-aln, bwa-mem*, the hybrid *bwa-aln-mem*, and *bowtie2*) using the same simulated samples as in section 3.2 (Table 1). For this comparison, all datasets were processed using the optimal mapping strategy established above (metagenomic filtering combined with competitive mapping) and default parameters (read merging enabled, minimum read length: 30 bp, mapping quality: 25). In sim-1, all mapping tools performed well, achieving 100% precision. However, the hybrid *bwa-aln-mem* approach and standard *bwa-mem* yielded the highest recall rates (90.57% and 90.54%, respectively) (Fig. 4). As sample degradation and contamination increased, stark performance differences emerged, highlighting a trade-off between precision and recall. In the highly contaminated sim-4 sample, *bwa-aln* successfully recovered the highest fraction of authentic ancient fragments (recall: 70.73%), but this sensitivity came at the cost of mapping accuracy; precision dropped to 75.98% due to the spurious alignment of 178 bacterial reads. The hybrid *bwa-aln-mem* strategy mirrored this exact behaviour in sim-4, maintaining the 70.73% recall rate but suffering the same drop in precision. Conversely, *bwa-mem* and *bowtie2* maintained 100% precision. This is because both of these tools do not allow enough mismatches in short reads, resulting in a noticeable penalty to sensitivity. In sim-4, *bwa-mem* and *bowtie2* recall rates dropped to 65.70% and 66.08%, respectively (Fig. 4). This indicates that these algorithms discarded true ultrashort endogenous fragments, which can create reference bias as reads with alternative alleles are preferentially rejected (as discussed in [9]). Together, these results demonstrate that no single mapping algorithm is universally optimal for all palaeogenomic contexts, due to the trade-off between precision and recall rates depending on contamination levels. The hybrid *bwa-aln-mem* approach is recommended for samples that contain a wide distribution of both short and long fragments and are mapped to a divergent reference genome. In such cases, the hybrid approach uniquely overcomes the length-dependent biases of using either algorithm alone (see Supplementary Text S2). By offering all four strategies, DNAharvester allows researchers to tailor the alignment process to the specific preservation state and fragment length distribution of their samples.

**Fig. 4.**
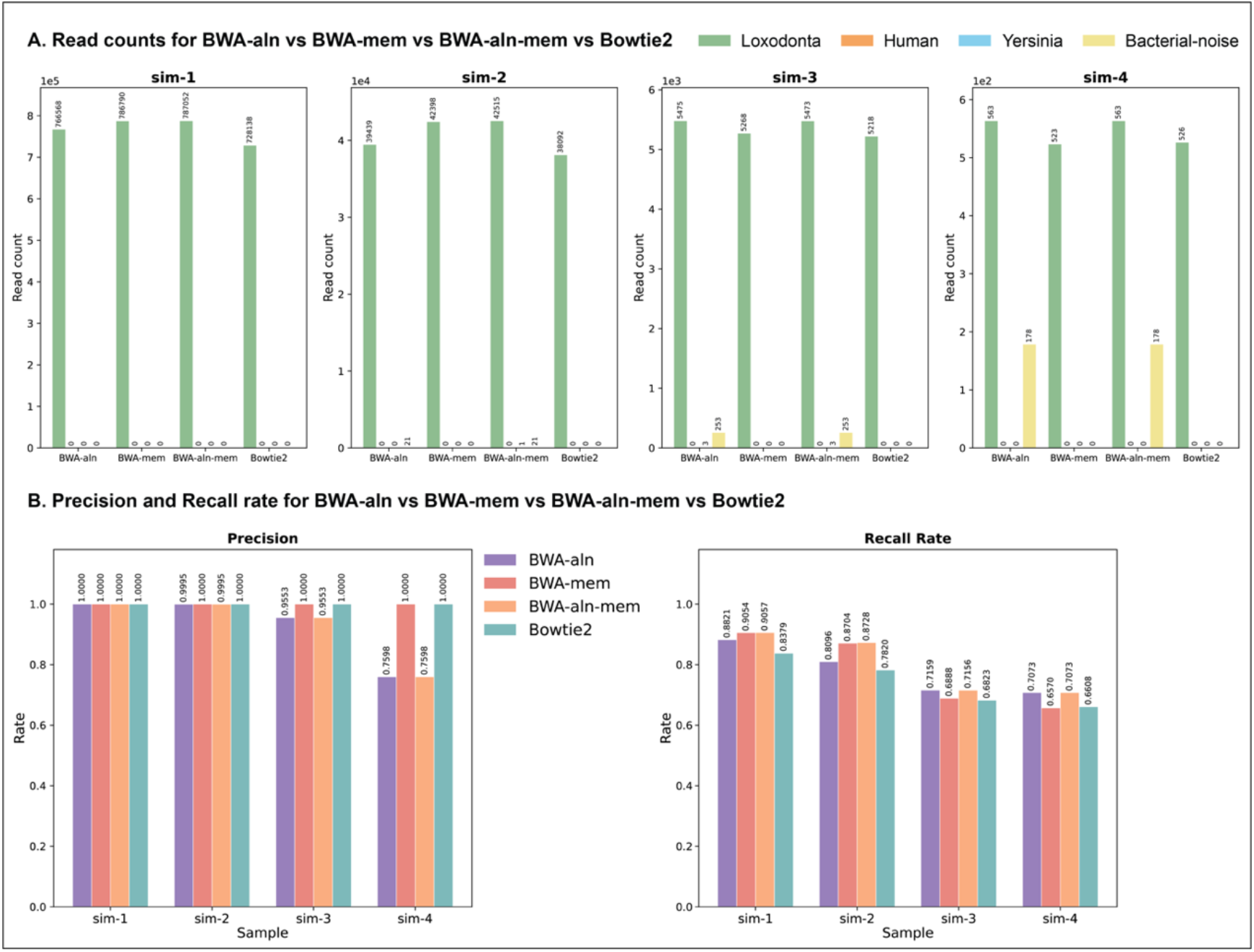
Performance comparison of different mapping tools available in DNAharvester. (A) Comparison of mapped read compositions resulting from the four supported mapping algorithms (*bwa-aln, bwa-mem, bwa-aln-mem*, and *bowtie2*). (B) The corresponding precision and recall metrics for each mapping tool. All datasets were processed using the optimal combined strategy of metagenomic pre-filtering and competitive mapping.

### 3.4. Iterative mitogenome assembly

To evaluate the iterative assembly subworkflow, varying numbers of simulated woolly mammoth mitochondrial reads were spiked into the simulated libraries at quantities of either 100, 1,000, 2,000, 5,000, or 10,000 reads for each of the three libraries for samples sim-1 through sim-5 (Table 2). These reads were also simulated using Gargammel [44] with lengths ranging from 30 to 100 bp, and post-mortem DNA damage patterns were applied using the Briggs model [8] (-damage 0.03,0.4,0.01,0.3). The mammoth mitogenome was then reconstructed using DNAharvester’s iterative assembly subworkflow with default settings (coverage_threshold = 3; call_fraction = 0.66; quality_threshold = 40), using the Asian elephant mitogenome (GenBank: CM123153.1) as the seed reference. The resulting consensus sequences (Supplementary Dataset S1) were aligned against the original mammoth mitogenome using MAFFT [45] and corrected for differing start and end coordinates using RefineAlign. As expected, the completeness of the iteratively assembled mitogenome scaled directly with the total number of endogenous reads available in the dataset (Table 2). In sim-1 (which contained 300 total spiked reads across three libraries), the pipeline recovered 10.49% of the mitochondrial genome. However, with 6,000 total spiked reads (sim-3), the pipeline successfully reconstructed 95.51% of the mitogenome sequence. At optimal coverage levels (sim-3 to sim-5), the reconstructed consensus sequences achieved near 100% accuracy when compared to the original mammoth reference. The few errors observed in sim-3 and sim-5 are located in the problematic VNTR (variable number of tandem repeats) section of the control region that is usually excluded from downstream analysis. Overall, these results confirm that the iterative assembly subworkflow can effectively and accurately reconstruct an extinct mitogenome utilising a closely related species as a seed reference.

### 3.5. Taxonomic classification

To evaluate the taxonomic classification subworkflow, we used the same simulated datasets described in section 3.4 (sim-1 through sim-5), which contain varying amounts of spiked simulated woolly mammoth mitochondrial reads. The analysis was performed using DNAharvester’s default taxonomic classification parameters (tc_max_alignments = 10; tc_mapping_quality = 1). For the reference database, we employed the complete RefSeq mitochondrial genome database (accessed on September 5, 2025: https://ftp.ncbi.nlm.nih.gov/refseq/release/mitochondrion/). The pipeline successfully identified the woolly mammoth mitogenome (NC_007596.2) as the top taxonomic hit across all samples (Table 3; Supplementary Dataset S1). The number of reads assigned to this top hit scaled proportionally with the total number of spiked reads. Notably, in sim-1, where a total of 300 mammoth mitochondrial reads were added, 324 reads were assigned to the mammoth mitogenome. These excess reads are due to the spurious mapping of African elephant autosomal reads, likely derived from nuclear mitochondrial DNA segments (NUMTs).

**Table 3.** Summary of taxonomic classification results. Identification of the top (woolly mammoth) and second (Columbian mammoth; *M. columbi*) taxonomic hit and assigned read counts using DNAharvester’s taxonomic classification subworkflow on five simulated samples (sim-1 to sim-5). The RefSeq mitochondrial database was used as the reference.

| Sample ID | Reads added per sample | Top hit | Top hit mapped reads | Second hit | Second hit mapped reads |
| --- | --- | --- | --- | --- | --- |
| sim-1 | 300 | NC_007596.2 | 324 | NC_015529.1 | 307 |
| sim-2 | 3,000 | NC_007596.2 | 2859 | NC_015529.1 | 2710 |
| sim-3 | 6,000 | NC_007596.2 | 5619 | NC_015529.1 | 5320 |
| sim-4 | 15,000 | NC_007596.2 | 13422 | NC_015529.1 | 12700 |
| sim-5 | 30,000 | NC_007596.2 | 25195 | NC_015529.1 | 23863 |

**Table 4.** Feature comparison between DNAharvester and existing ancient DNA pipelines.

| Feature | DNAharvester | GenErode | EAGER | PALEOMIX |
| --- | --- | --- | --- | --- |
| FastQ processing (QC, adapter trimming, read merging) | ✓ | ✓ | ✓ | ✓ |
| Metagenomic pre-filtering | ✓ | ✗ | ✗ | ✗ |
| Mapping to the target genome | ✓ | ✓ | ✓ | ✓ |
| Competitive mapping | ✓ | ✗ | ✗ | ✗ |
| Auto-estimation of read length cutoff | ✓ | ✗ | ✗ | ✗ |
| Damage profiling | ✓ | ✓ | ✓ | ✓ |
| Library complexity estimation | ✓ | ✗ | ✓ | ✗ |
| Contamination estimation | ✗ | ✗ | ✓ | ✗ |
| Repeat identification | ✓ | ✓ | ✗ | ✗ |
| CpG site identification | ✓ | ✓ | ✗ | ✗ |
| Variant calling | ✓ | ✓ | ✓ | ✓ |
| Genotype likelihood / pseudo-haploid calling | ✓ | ✗ | ✓ | ✗ |
| Iterative assembly of the mitogenome | ✓ | ✗ | ✗ | ✗ |
| Nuclear consensus sequence generation | ✗ | ✗ | ✓ | ✓ |
| Genetic sex determination | ✓ | ✗ | ✓ | ✗ |
| Taxonomic classification | ✓ | ✗ | ✓ | ✗ |
| Microbial screening | ✓ | ✗ | ✓ | ✗ |
| Phylogenetic analysis | ✗ | ✗ | ✗ | ✓ |
| Population genetic analyses (heterozygosity, ROH, PCA) | ✗ | ✓ | ✗ | ✗ |
| SNP functional annotation | ✗ | ✓ | ✗ | ✗ |
| Mutational load estimation | ✗ | ✓ | ✗ | ✗ |

### 3.6. Microbial screening

The microbial screening subworkflow was benchmarked using the simulated datasets described in Table 1. The data were processed using the default settings of DNAharvester’s microbial screening subworkflow (ms_mapping_tool = *bwa-aln*; ms_mapping_quality = 1). For this analysis, we used a custom reference database composed of the default pathogen list from Heuristic Operations for Pathogen Screening (HOPS), supplemented with additional animal-specific genomes (provided in Supplementary Dataset S1). Additionally, a novel custom Python script (pathogen_reference_database_update.py) is provided alongside the pipeline code, allowing users to easily generate custom databases or append new genomes to an existing database using a list of NCBI accession IDs. The microbial screening subworkflow successfully detected the presence of *Yersinia pestis* as a potential target in all simulated samples, with the exception of sim-5, which served as a negative control. In the sim-4 sample, *Yersinia pestis* constituted only 0.1% of the total reads (n = 999) (Table 1, Fig. 2A). Out of these, 137 reads were successfully recovered and exhibited the ancient DNA damage patterns expected from the simulated parameters. All generated PDF report files are provided in Supplementary Dataset S1.

### 3.7. Genetic sex determination

To validate the sexing subworkflow, we used previously published sequencing data from 11 human samples representing five different karyotypes (XX, XY, XXY, XYY, and X0) [43]. A random subset of one million reads per sample was mapped using DNAharvester’s default settings to the same reference genome as the original study (hs37d5), and all three genetic sex determination methods supported by the pipeline were applied. Consistent with the original study [43], DNAharvester accurately classified all five karyotypes (Fig. 5, Supplementary Dataset S1).

**Fig. 5.**
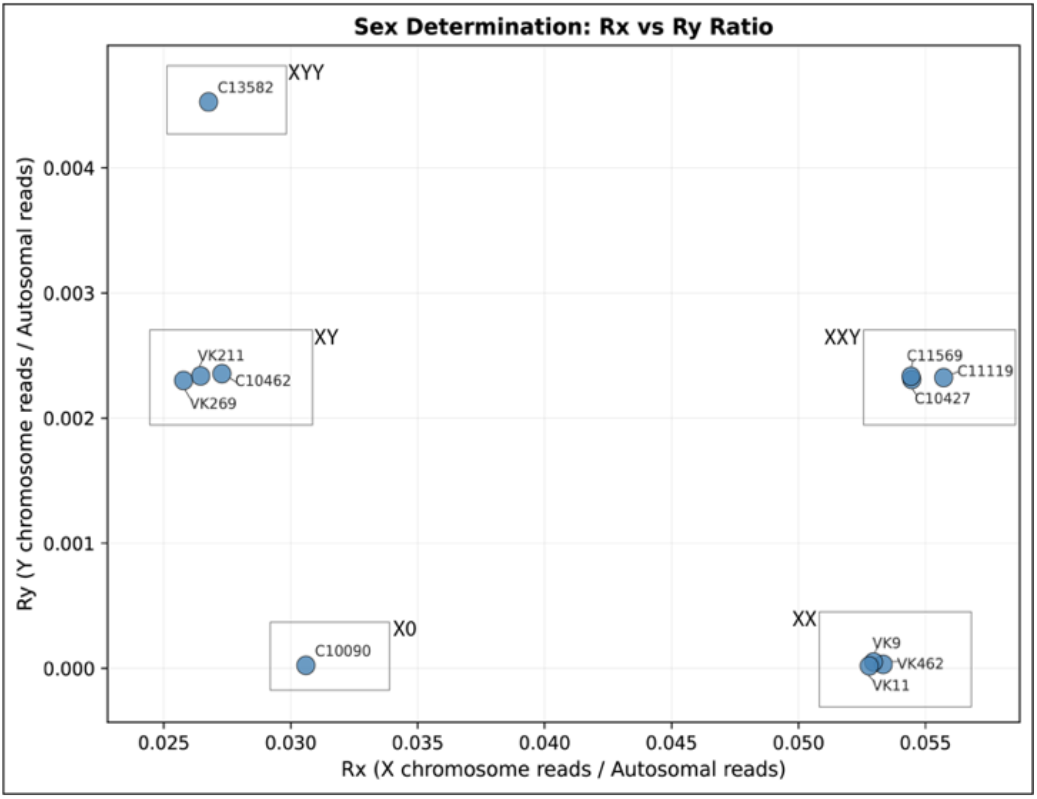
Genetic sex determination using DNAharvester. Scatter plot of normalised sex chromosome read depths comparing the X/Autosome (Rx) ratio against the Y/Autosome (Ry) ratio for 11 previously published human samples representing five distinct karyotypes (XX, XY, XXY, XYY, and X0) [43]. The boxes with karyotype labels were added manually for visualisation purposes after the DNAharvester run.

### 3.8. Computational requirements

To benchmark the computational requirements of DNAharvester, we generated simulated datasets containing 1, 10, 100, and 300 million read pairs per sample, each comprising the same four samples described in section 3.1 (sim-1 to sim-4) with the compositions, fragment length distributions, and damage parameters given in Table 1. All other parameters were held constant, including the reference genome, the number of samples and libraries, the workflow steps executed, and the CPU allocation of every process. Runs were performed on Dardel HPC compute nodes, each with two AMD EPYC− Zen2 2.25 GHz 64-core processors, and per-task CPU and memory consumption were recorded using Nextflow’s execution tracing.

Total CPU time scaled linearly with input size, with mapping dominating the computational cost across all dataset sizes (accounting for 90.1% to 99.2% of the total; Fig. 6A-B, Table S1). In contrast, peak memory scaled modestly, increasing only 5.1-fold (from 15.3 GB to 78.8 GB) across a 300-fold increase in input reads (Fig. 6C). For datasets of up to 100 million read pairs per sample, BWA indexing and alignment sorting buffers were the most memory-intensive components. However, for the largest datasets, fastp overtook mapping as the primary memory consumer. As a practical reference point, processing 100 million read pairs against the 3.4 Gb Asian elephant reference genome required approximately 282 CPU-hours. Here, GPU-accelerated short-read aligners represent a promising avenue for further reducing processing time.

**Fig. 6.**
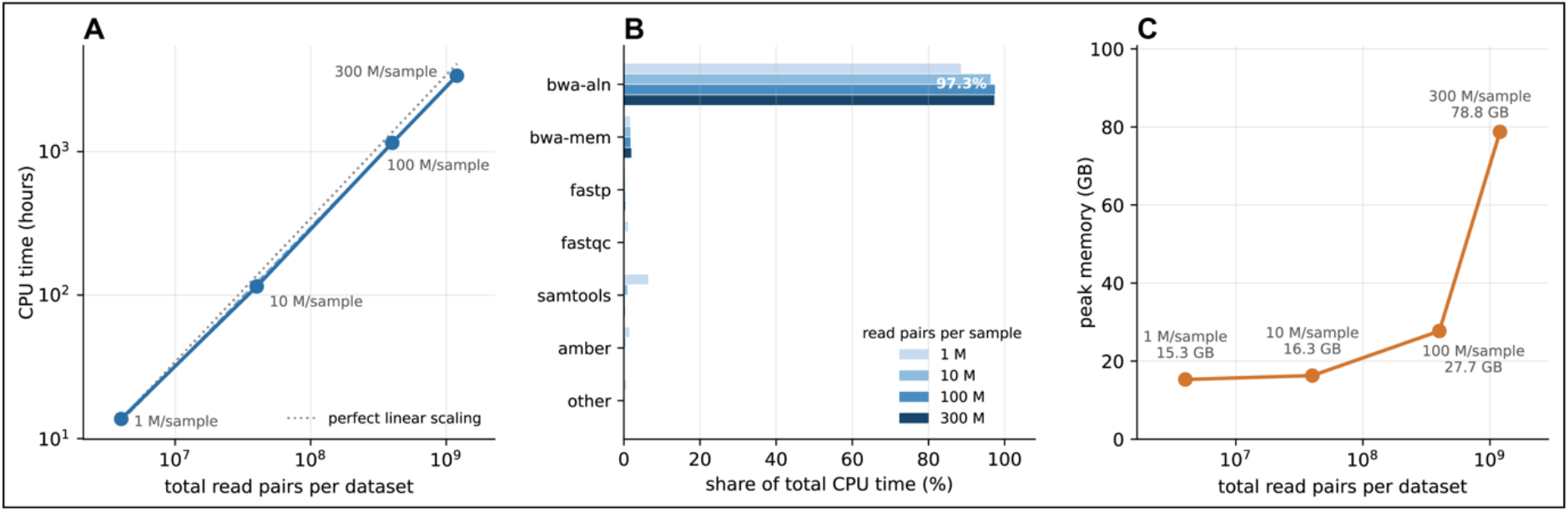
Computational resource requirements of DNAharvester. Four simulated datasets, differing only in read pairs per library, processed with an identical configuration. Points and bars are labelled by read pairs per sample. **(A)** Total CPU time versus dataset size. **(B)** Share of total CPU time by process. **(C)** Peak memory usage.

## 4. Accessibility, Reproducibility, and Scalability

### 4.1. Accessibility and documentation

DNAharvester is designed to be user-friendly, requiring only basic command-line experience, while still allowing for flexibility in the analyses for experienced users. It includes detailed documentation covering all aspects of the pipeline, from input data preparation to result interpretation. Users provide a sample sheet listing their sequencing data and a configuration file to customise workflow parameters. This streamlined setup ensures accessibility for researchers across disciplines, facilitating the standardisation of ancient DNA data processing. The source code, documentation, and example datasets are available on GitHub at https://github.com/NBISweden/DNAharvester.

### 4.2. Reproducibility

Implemented in Nextflow [46], DNAharvester is designed in adherence to the FAIR data principles (Findable, Accessible, Interoperable, and Reusable), providing reproducibility and portability across different computing environments. The pipeline supports multiple software management systems, including Conda [47] and Mamba [48], as well as containerization platforms including Docker [49], Apptainer [50], and Singularity [50]. By providing software dependencies in containers, DNAharvester eliminates version conflicts and guarantees that analyses can be replicated exactly, regardless of the underlying infrastructure.

### 4.3. Scalability and execution

DNAharvester is highly scalable, capable of processing datasets ranging from small pilot studies to large-scale genomic projects. As demonstrated in our benchmarking (Section 3.8), computational time scaled linearly with read count, driven primarily by bwa-aln mapping, while peak memory grew 5.1-fold across a 300-fold increase in data volume. The pipeline can be executed on a local machine for smaller datasets (e.g., 5-10 samples, with 10-20 million reads per sample) or deployed on High-Performance Computing (HPC) clusters for computationally intensive tasks. DNAharvester includes pre-configured profiles for common workload managers like SLURM [51], allowing efficient resource allocation and parallel processing.

## 5. Summary

DNAharvester is a comprehensive and modular Nextflow pipeline tailored specifically for highly degraded and complex palaeogenomics datasets. While existing workflows, such as GenErode [33], EAGER [52], and PALEOMIX [53], provide essential standard processing steps like fastq processing, mapping, and variant calling, DNAharvester distinguishes itself by integrating novel and state-of-the-art features designed specifically for ancient samples with low endogenous content. The pipeline fully automates metagenomic filtering and competitive mapping, drastically reducing noise from spurious alignments. It also supports multiple mapping tools and dynamic auto-estimation of read length cutoffs. Furthermore, the pipeline offers subworkflows for automated iterative assembly of mitogenomes, taxonomic classification, genomic repeats and CpG sites identification, multiple variant calling options for low to high coverage samples, genetic sex determination, and microbial screening for pathogen detection. Benchmarking across a 300-fold range in dataset size confirmed that its computational cost scales linearly with read count, while peak memory grows 5.1-fold over the same range.

Importantly, unlike some of its counterparts, DNAharvester does not perform downstream population-genetic analyses. Instead, its true power lies in rigorous and flexible upstream data processing. By providing advanced filtering and mapping strategies within a fully containerised and scalable framework, the pipeline ensures the maximum recovery of authentic endogenous ancient DNA data while systematically mitigating background noise.

## Acknowledgments

The work was funded by the European Research Council (Primigenomes, 101054984, awarded to LD). VEK was supported by the SciLifeLab & Wallenberg Data Driven Life Science Program, Knut and Alice Wallenberg Foundation (grants: KAW 2020.0239 and KAW 2017.0003), and by the National Bioinformatics Infrastructure Sweden (NBIS) at SciLifeLab. NO is supported by the European Regional Development Fund project “Functional Omics Analysis of Metabolic Diseases to Advance Drug Discovery Research Excellence in LIOS - TARGETWISE” (No.1.1.1.5./2/24/A/003). The authors also acknowledge support from Knut and Alice Wallenberg Foundation grants KAW 2021.0048 (PDH) and KAW 2022.0033 (PDH and LD), as well as from Swedish Research Council grant 2021-00625 (LD). We are grateful to all the staff/students at the Centre for Palaeogenetics (CPG) for their valuable discussions during the development of the pipeline. The development of the pipeline was enabled by computational resources provided by the National Academic Infrastructure for Supercomputing in Sweden (NAISS), partially funded by the Swedish Research Council through grant agreement no. 2022-06725 (project id: NAISS 2025/5-78 and NAISS 2025/22-1155). During the preparation of this manuscript, a large language model (Google Gemini) was used to improve grammar and sentence structure, and the authors take full responsibility for the final text.

## Author Contributions

Conceptualisation and Methodology: All authors

Pipeline development: BS, VEK

Pipeline testing and Review: BS, VEK

Pipeline benchmarking: BS

Funding acquisition: LD

Project administration: LD

Supervision: LD, PDH, NO, DDM

Visualisation: BS, VEK

Writing – original draft: BS

Writing – review & editing: All authors

## Data and Code Availability

All genomes used for the simulated datasets are publicly available at the RefSeq accessions given in the text or the GTDB. The source code, documentation, and example datasets are available on GitHub at https://github.com/NBISweden/DNAharvester, under the GNU General Public License v3.0 (GPLv3) All scripts used to generate the simulated datasets (sim-1 to sim-5), the Nextflow configuration files used for benchmarking, and the code required to reproduce the analyses and figures presented in this manuscript are available in the GitHub repository: https://github.com/bilalbioinfo/DNAharvester-Sharif-et-al-2026. The RefineAlign script for generating the mitogenome alignment is available at the GitHub repository: https://github.com/pheintzman/RefineAlign. The result files generated during the pipeline benchmarking, including bam, fasta, idxstats, and pdf files (Supplementary Dataset S1), are publicly available on Zenodo repository: https://doi.org/10.5281/zenodo.22725211.

## Supplementary Text

The Supplementary Text, including supplementary figures, is available as an attached file alongside this manuscript, and a copy is also included with Supplementary Dataset S1 above.

