## Supplementary Text for "DNAharvester: A Nextflow Pipeline for Analysing Highly Degraded DNA from Ancient and Historical Specimens"

#### Text S1. Additional adapter removal with AdaptClean

Fastp [12] requires a minimum sequence overlap of 5 bp in order to be able to identify an adapter sequence. This safeguards against trimming insert sequences, but it creates a blind spot for short adapter remnants (of 1–4 bp at read ends). For example, sequences like A, AG, or AGAT often remain attached to the 3' end of the read because they fall below the detection threshold. This creates a specific artifact in the data: the processed fastq file will be missing reads with lengths immediately below the maximum sequencing cycle length (defined here as $L_{max}$). Consequently, reads ranging from ($L_{max}$ – 1) to ($L_{max}$ – 4) are effectively absent. These untrimmed adapter bases are problematic because they introduce erroneous sequences to the ends of the reads. During mapping, this increases mismatch rates and can cause reference bias. To demonstrate this issue, we simulated 1 million single-end reads using Gargammel [41] (-n 1000000 --comp 0,0.5,0.5 --loc 4.2 --scale 0.6 --minsize 20 --maxsize 100 -rl 100 -se -damage 0,0,0,0 -qs 93), comprising 50% reads from African elephant (*Loxodonta africana*) chromosome 1 (GenBank: NC_087342.1) and 50% from human (*Homo sapiens*) chromosome 1 (GenBank: NC_000001.11). The simulated reads were mapped to NC_087342.1, using DNAharvester's default settings, except with and without AdaptClean. When using only Fastp, read lengths from 96 to 99 bp were completely absent, and a higher mismatch rate was observed for reads of exactly 100 bp (Fig. S1). To fix this issue, we implemented AdaptClean [13] in DNAharvester, a custom C++ script that specifically targets only those reads that have not been trimmed (*i.e.*, reads equal to the full sequencing cycle length). It removes the specific 1–4 bp adapter remnants from these reads without affecting the rest of the dataset. This recovers the missing read lengths and ensures that the ends of the reads are free of technical artefacts (Fig. S1).

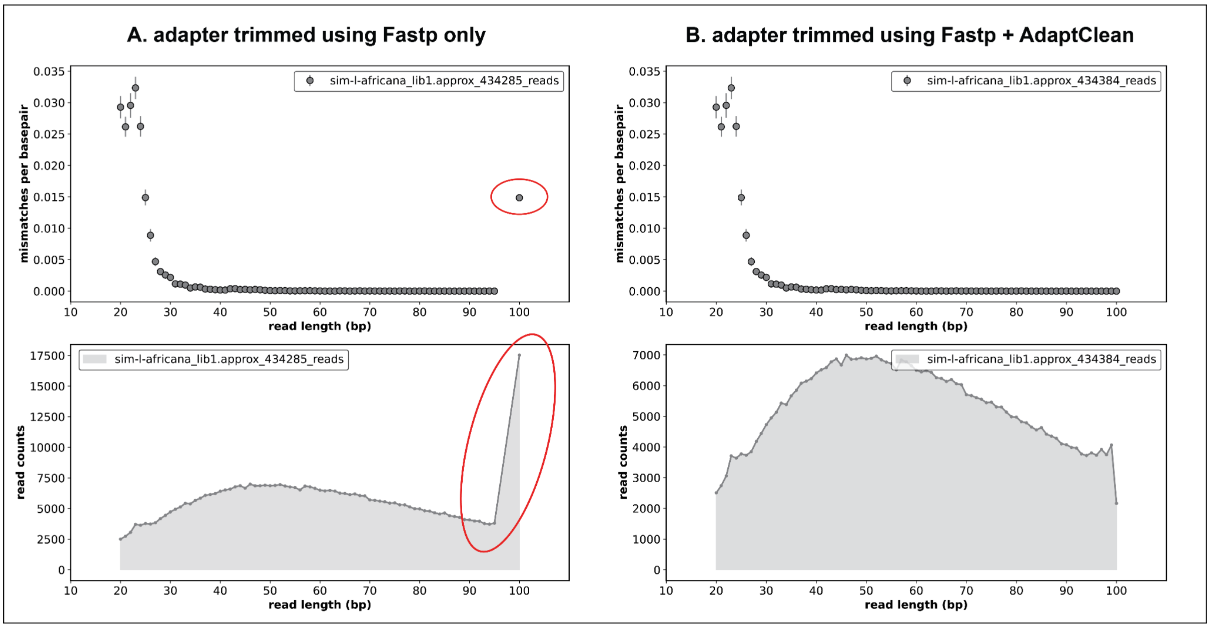


**Fig. S1.** AMBER plot illustrating the disparity in mismatch rate and read length distribution (highlighted by the red ellipses) when comparing adapter removal solely with Fastp (A) versus using the combination of Fastp and AdaptClean (B).

### Text S2. The hybrid *bwa-aln-mem* mapping strategy

A hybrid alignment tool, *bwa-aln-mem*, is included in DNAharvester to accommodate samples containing both short and long DNA fragments. This approach uses *bwa-aln* for aligning short reads and *bwa-mem* for reads exceeding a predefined length threshold (default: 70 bp). To demonstrate its need, here, we simulated 1 million reads from Lion (*Panthera leo*) chromosome 1 (GenBank: CM101310) using Gargammel [41], and mapped them to the domestic cat reference assembly (GCA_051911835.1), representing a sequence divergence of approximately 2%. The reads were mapped using both *bwa-aln* and *bwa-mem*, and a combined AMBER plot was generated (Fig. S2). For reads between 30 and ~80 bp, *bwa-aln* successfully recovers the expected 2% divergence. However, the mismatch rate artificially drops for longer reads because *bwa-aln* restricts the maximum number of allowed mismatches. Conversely, for reads shorter than ~80 bp, *bwa-mem* exhibits strong reference bias by failing to map short, divergent reads. However, *bwa-mem* correctly captures the expected mismatch rate for longer fragments. This aligner-specific bias becomes even more severe in datasets with higher divergence, such as a simulated sample where we artificially introduced a 4% sequence divergence to 1 million reads generated from Asian elephant (*Elephas maximus*) chromosome 26 (GenBank: NC_064844.1) and mapped back to the same chromosome (Fig. S3). To capitalise on the strengths of both algorithms in DNAharvester, we implement a hybrid mapping tool that automatically splits the input FASTQ files into short- and long-read fractions, aligns each fraction with its optimal algorithm, and merges the resulting BAM files. Ultimately, this combined *bwa-aln-mem* mapping tool is highly advantageous when aligning ancient or degraded samples to highly divergent reference genomes.

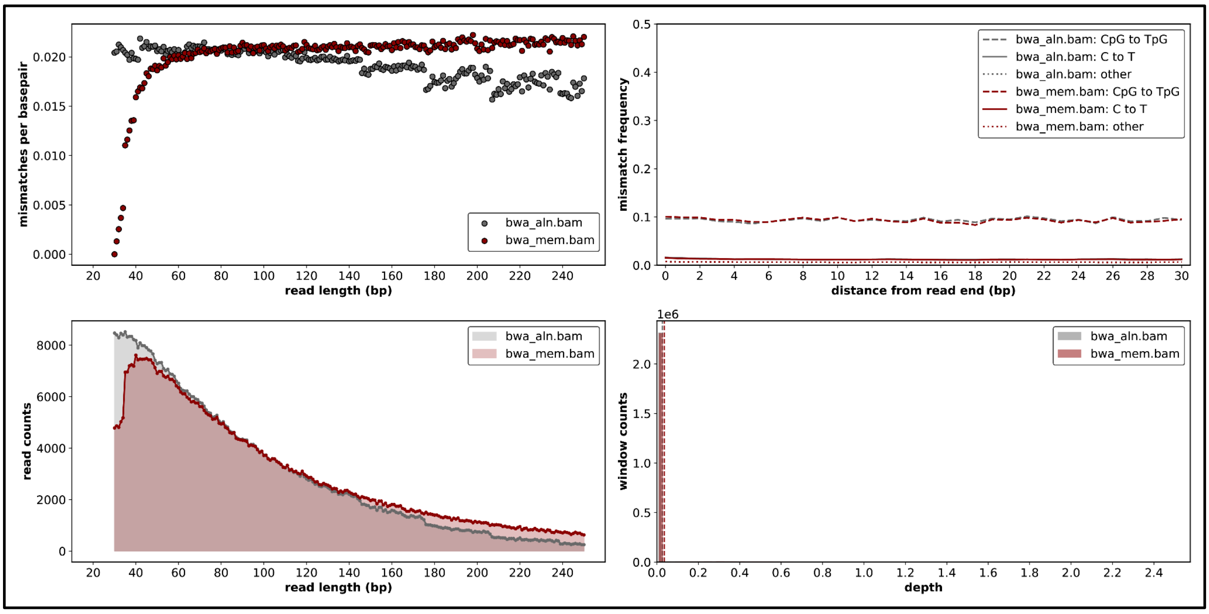


**Fig. S2. Impact of read length and aligner choice on reference bias at ~2% sequence to reference divergence.** AMBER plot illustrating the mismatch rates across varying read lengths for 1 million simulated reads from Lion (*Panthera leo*) chromosome 1 (CM101310) mapped to the domestic cat reference assembly (GCA_051911835.1).

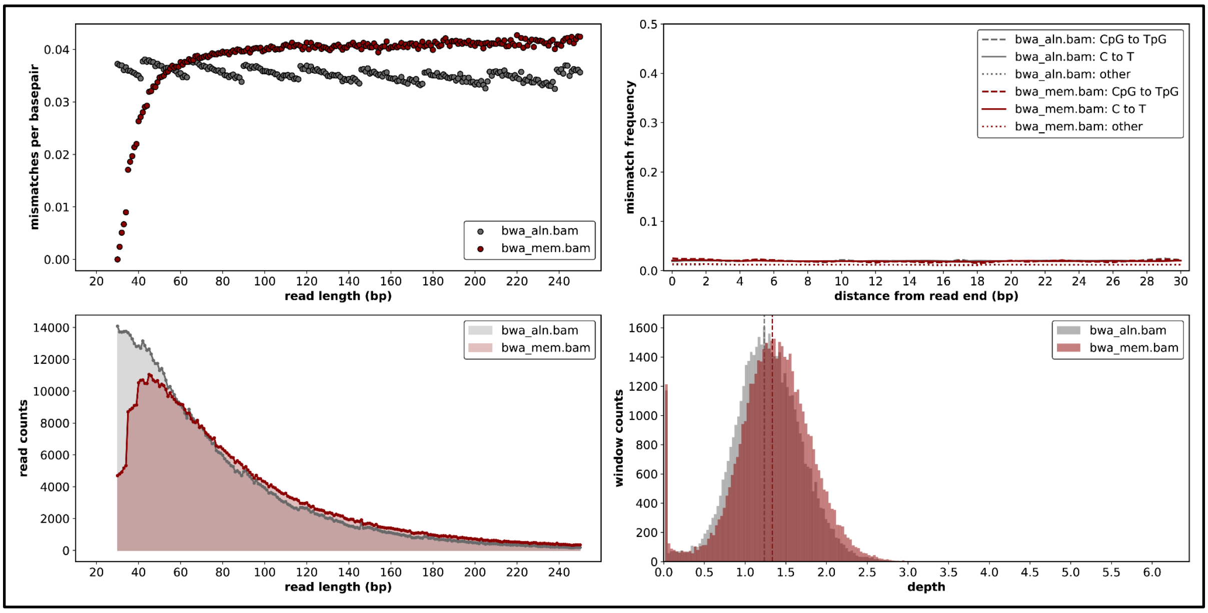


**Fig. S3. Impact of read length and aligner choice on reference bias at 4% sequence to reference divergence.** AMBER plot illustrating the mismatch rates across varying read lengths for 1 million simulated reads from Asian elephant (*Elephas maximus*) chromosome 26 (NC_064844.1) with an artificially introduced 4% sequence divergence and mapped to NC_064844.1.

### Text S3. Automatic minimum read length cutoff

Theoretically, the mismatch rate between ancient DNA molecules and the reference genome should remain constant across different read lengths. However, in reality the mismatch rate in ultrashort reads (typically <35 bp) often deviates from this baseline, especially if endogenous content is low [9]. This deviation manifests differently depending on the mapping algorithm: *bwa-aln* typically exhibits an increased mismatch rate for ultrashort reads due to the accumulation of spurious alignments, whereas *bwa-mem* shows a decreased mismatch rate driven by strong reference bias. To systematically filter these problematic reads without arbitrarily discarding valid data, DNAharvester dynamically estimates a minimum read length cutoff using the Knee Point Detection algorithm (kneed). The pipeline parses AMBER mismatch-rate metrics for fragments under 40 bp. To ensure statistical robustness, a minimum of 10,000 mapped reads is required to model the curve; samples falling below this depth default to a 30 bp cutoff. For qualifying datasets, a custom Python script identifies the exact inflection point (the "knee") where the error rate diverges from the stable baseline, dynamically adjusting its curve-fitting parameters (convex or concave) based on the specific aligner used. The definitive read length cutoff is then set to the knee point plus one base, effectively removing erroneous ultrashort fragments while seamlessly maximizing the retention of authentic ancient DNA data.

To illustrate this, we processed published data (20 million reads) from the 1.25-million-year-old mammoth "Adycha" [6] and mapped the reads to the Asian elephant (*Elephas maximus*) reference genome (GCF_024166365.1), alongside the decoy human genome (GCF_000001405.40). This was performed using the default settings in DNAharvester (*bwa-aln*, read merging enabled, minimum read length: 30 bp, mapping quality: 25), except the read length parameter was set to auto. In this example, the optimal read length cutoff was found to be 30 bp (Fig. S4), and all reads below this threshold were automatically discarded in the subsequent processing step (see Fig. 1).

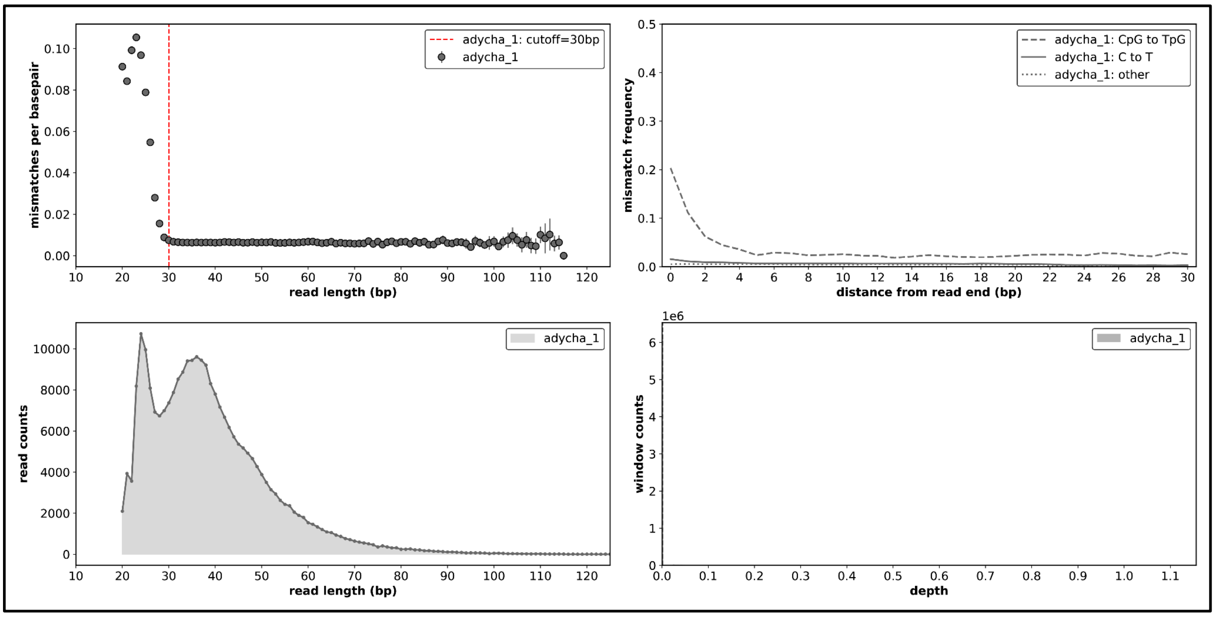


**Fig. S4.** Automated read length cutoff estimation. AMBER plot illustrating the mismatch rate per read length for published data from the 1.25-million-year-old mammoth "Adycha" mapped to the Asian elephant (*Elephas maximus*) reference genome using *bwa-aln*. The vertical red line indicates the optimal minimum read length cutoff (calculated here as 30 bp) dynamically determined by DNAharvester to filter out spurious alignments. (Note: DNAharvester uses the standard, published version of AMBER [9], the figure was generated using a custom-modified version exclusively for this manuscript to visualize the red cutoff line.)

### Text S4. GATK IndelRealignment

When mapping short ancient DNA fragments to a highly divergent reference genome, standard aligners often struggle to correctly position insertions and deletions (indels). Specifically, because BWA-aln uses significantly higher scoring penalties for gap openings (indels) compared to base mismatches (SNVs), it may force mismatched bases instead of appropriately opening a gap. This results in clusters of false single-nucleotide variants, particularly near the ends of reads. To mitigate this issue, DNAharvester incorporates an optional local realignment step using GATK3. This process evaluates all reads spanning an indel and realigns them jointly to minimize the total number of mismatching bases. As shown in Fig. S5, raw BAM without realignment often exhibit these spurious variants at the read termini, which GATK IndelRealignment effectively resolves.

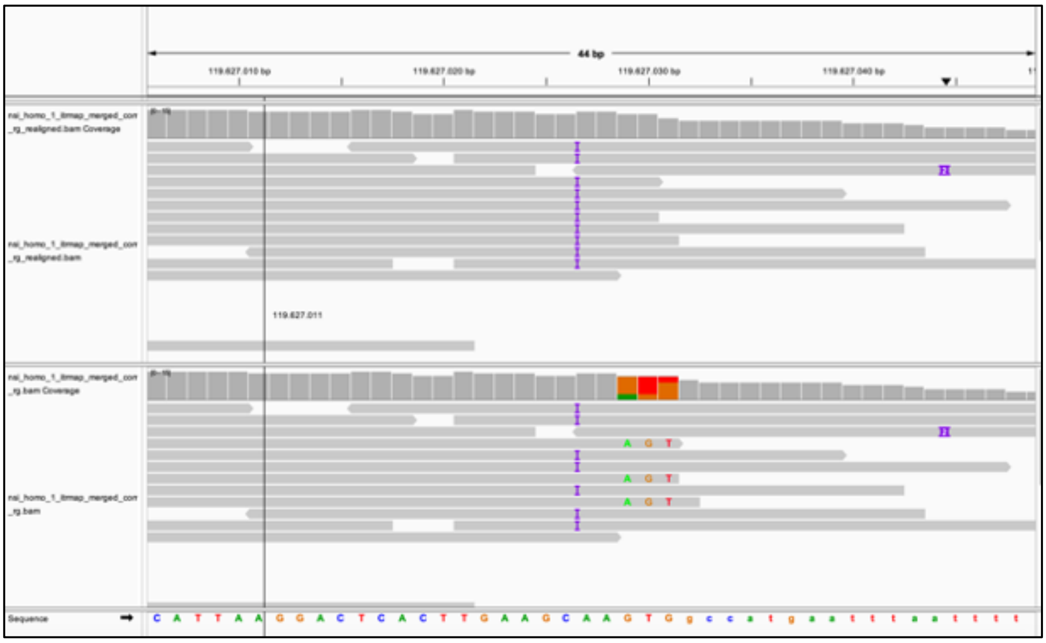

**Fig. S5. Impact of GATK IndelRealignment on misalignment artefacts.** Integrative Genomics Viewer (IGV) screenshots comparing read alignments. (Top) Alignments processed with GATK IndelRealignment. (Bottom) Default BWA alignments without realignment. In the un-realigned dataset, the aligner introduces false single-nucleotide variants at the read ends.

**Table S1. Computational resource requirements of DNAharvester.** Resource consumption for four simulated datasets of increasing size, each comprising four samples with three libraries per sample and processed using an identical configuration. CPU time is the sum across all tasks of task runtime multiplied by measured CPU utilisation; peak memory is the highest resident set size recorded for any single task.

| Read pairs per sample | Total read pairs per dataset | CPU time (h) | Peak memory (GB) |
|---|---|---|---|
| 1,000,000 | 3,999,987 | 13.7 | 15.3 |
| 10,000,000 | 40,000,065 | 114.9 | 16.3 |
| 100,000,000 | 400,000,779 | 1,152.2 | 27.7 |
| 300,000,000 | 1,200,002,346 | 3,386.3 | 78.8 |
